# MetaCerberus: distributed highly parallelized scalable HMM-based implementation for robust functional annotation across the tree of life

**DOI:** 10.1101/2023.08.10.552700

**Authors:** Jose L. Figueroa, Eliza Dhungel, Cory R. Brouwer, Richard Allen White

## Abstract

**Summary:** MetaCerberus is an exclusive HMM/HMMER-based tool that is massively parallel, on low memory, and provides rapid scalable annotation for functional gene inference across genomes to metacommunities. It provides robust enumeration of functional genes and pathways across many current public databases including KEGG (KO), COGs, CAZy, FOAM, and viral specific databases (i.e., VOGs and PHROGs). In a direct comparison, MetaCerberus was twice as fast as EggNOG-Mapper, and produced better annotation of viruses, phages, and archaeal viruses than DRAM, PROKKA, or InterProScan. MetaCerberus annotates more KOs across domains when compared to DRAM, with a 186x smaller database and a third less memory. MetaCerberus is fully integrated with differential statistical tools (i.e., DESeq2 and edgeR), pathway enrichment (GAGE R), and Pathview R for quantitative elucidation of metabolic pathways. MetaCerberus implements the key to unlocking the biosphere across the tree of life at scale.

**Availability and implementation:** MetaCerberus is written in Python and distributed under a BSD-3 license. The source code of MetaCerberus is freely available at https://github.com/raw-lab/metacerberus. Written in python 3 for both Linux and Mac OS X. MetaCerberus can also be easily installed using mamba create –n metacerberus –c bioconda –c conda-forge metacerberus

## Introduction

Annotation is a fundamental step in functional gene inference, which is required by many disciplines in biology. Massively parallel sequencing (MPS) has reached the terabyte scale with Illumina NovaSeq X producing 16 Tb per run and Oxford nanopore promethION 7 Tb per run (**1-2**). Due to this increase in MPS, the number of reference microbial genomes and metagenomes has increased by orders of magnitude. Genome Taxonomy Database (GTDB) now includes 402,709 (08-RS214, April 28^th^, 2023) genomes, and the Short Read Archive (SRA) has >4.5 million listed biosample metagenomes (**3-4**). Cellular metagenome-assembled genomes (MAGs) and their viral counterpart vMAGs (viral MAGs) have also rapidly populated public databases through reconstruction from shotgun metagenomics (**5-7**). Functional gene annotation is required for metabolic reconstruction, functional and structural gene differential analysis, inference of pathway regulation, presence/absence of toxin genes (e.g., botulinum toxin A), novel gene cluster discovery (e.g., antibiotic discovery), and viral detection. Due to this Terabyte scale, the annotation step will be the most prolonged, requiring more CPU time, memory, and resources to finish before obtaining biological insight. Reference databases have also been nearing the Terabyte scale, taking days to download and format, requiring massive allocations of disk space. Thus scalable, highly parallel, low memory, and rapid annotation tools are critical to the future of ‘omics analysis.

Functional annotation requires two main steps: 1) gene calling followed by 2) gene assignment via external reference databases. Multiple approaches have been applied for gene calling and gene assignment, including homology and ontology-based methods. Gene calling finds protein-coding open reading frames (pORFs) alongside ribosomal RNAs, transfer RNAs, and other RNAs. Various tools exist for pORF calling, including Prodigal, FragGeneScanRs, GetOrf, and GeneMark (**8-11**). Gene assignment of pORFs to external databases often uses homology-based tools such as BLAST (**12**), MMseq2 (**13**), and/or DIAMOND (**14**) against databases such as RefSeq (NCBI Reference Sequence Database) (**15**), UniProt (Universal Protein Resource) (**16**), or KEGG (Kyoto Encyclopedia of Genes and Genomes) (**17**). Common tools include PROKKA, DRAM (Distilled and Refined Annotation of Metabolism), InterProScan (INTEgrative PROtein signature database), EggNOG-Mapper (evolutionary genealogy of genes: Non-supervised Orthologous Groups), and MicrobeAnnotator (**18-22**). Ontology-based approaches are generally superior to homology-based methods (**21**). EggNOG-Mapper and InterProScan utilize homology-based (i.e., Diamond and BLAST) and Hidden Markov Models (HMMs) based ontology approaches via HMMER (**23**) using either KEGG (**17**), eggNOG (**24**), InterPro (**25**), or Pfam (protein family) databases (**26**). HMMs provide greater sensitivity to elucidate and discover relationships between query and database based on ontology and are protein domain-centric (**21, 27**).

Viruses and the candidate phyla radiation (CPR) have remained challenging to functionally annotate due to the divergent nature of their proteins (**28-29**). DRAM has a specific version (i.e., DRAM-v) to annotate viruses, including the detection of viral auxiliary metabolic genes (vAMGs) (**19**). MicrobeAnnotator and DRAM have attempted to close the gap in CPR functional annotation. While no specific annotation tool or gene database exists for CPR, they are found amongst GTDB and other public repositories (**30**). Various databases such as VOGs (Virus Orthologous Groups), pVOGs (Prokaryotic Virus Orthologous Groups), IMG/VR (Integrated Microbial Genome/Virus), INPHARED (INfrastructure for a PHAge REference Database), and PHROGs (Prokaryotic Virus Remote Homologous Groups database) have been introduced to improve annotation viruses from isolates and vMAGs (**31-35**). Still, CPR and viruses remain a significant challenge for functional annotation.

Many tools are avaliable for functional annotation from genomes to metagenomes; however, gaps remain between: 1) resource utilization (e.g., memory use), 2) large database size, and 3) parallel processing, and simultaneously providing robust rapid annotation at scale. Further development of tools for CPR and viral functional annotation are a general community need. We present MetaCerberus, an ontology-based HMM tool that provides scalable, highly parallel, low memory usage, and rapid annotation for genomes to metacommunities across the tree of life.

## Implementation

### Framework and coding base

MetaCerberus is written entirely in Python (version 3) as a wrapper for various other tools described below. Similar to our other software MerCat2 for massively parallel processing (MPP), it utilizes a byte chunking algorithm 1 (’Chunker’) to split files for MPP for further utilization in RAY, a massive open-source parallel computing framework to scale Python applications and workflows (**36**). Using RAY’s scalable parallelization within MetaCerberus allows utilization across multiple nodes with ease. To avoid large RAM consumption, we implemented the greedy algorithm for tab-separated merging and incremental PCA plot limiter from MerCat2 (**36**). MetaCerberus utilizes HMM/HMMER exclusively without homology-based tools (e.g., BLAST)

### Databases for MetaCerberus

MetaCerberus enables functional gene assignment across multiple databases, including: 1) KOfams (KEGG protein families) to obtain KEGG KOs (KEGG Ontology) (version 11-Jul-2023, https://www.genome.jp/ftp/db/kofam/), 2) FOAM (Functional Ontology Assignments for Metagenomes), 3) COG (Clusters of Orthologous Genes) (version 2020, https://ftp.ncbi.nih.gov/pub/COG/COG2020/data/), and 4) dbCAN (DataBase for automated Carbohydrate-active enzyme ANnotation) for CAZy (Carbohydrate-Active enZYmes Database) (version 11, https://bcb.unl.edu/dbCAN2/download/) (**37-41**). For viral annotation, MetaCerberus enlists VOG (version 219, https://vogdb.org/download), pVOG (version Sep2016, https://ftp.ncbi.nlm.nih.gov/pub/kristensen/pVOGs/downloads.html#), and PHROG (version 4, https://phrogs.lmge.uca.fr/) databases. FOAM ontology is obtained from KOfam KOs, and then computed via a reference table to avoid redundancy. Similarly, the dbCAN database is used to obtain CAZy ontology via a reference table. COGs and PHROGs are currently not formatted as HMMs within their public repositories. We converted them into protein family-specific HMMs (e.g., COG1 –> COG1.hmm) using MAFFT (version 7.273-woe) via local alignments with maximum iterations of 1000 (**42**). We compared databases of six other tools to MetaCerberus, including DRAM, PROKKA, InterProScan, MicrobeAnnotator, and EggNOG-Mapper (**Table 1**). Currently, only MetaCerberus provides functional annotation and support to FOAM, pVOG, and PHROG databases (**Table 1**). EggNOG-Mapper and MetaCerberus are the only tools we compared that supports the COG database (**Table 1**). All tools compared in this study obtain the enzyme commission numbers (EC) numbers (**Table 1**).

**Table 1:** Comparing tools based on databases provided. This includes versions of other databases present within the various tools compared.

|  | EC | KEGG | CAZy | COG | FOAM | VOG | pVOG | PHROG | pfam | EggNOG | InterPro |
| --- | --- | --- | --- | --- | --- | --- | --- | --- | --- | --- | --- |
| MetaCerberus | X | X | X | X | X | X | X | X |  |  |  |
| DRAM | X | X | X |  |  | X |  |  | X |  |  |
| Prokka | X |  |  |  |  |  |  |  | X |  |  |
| InterProScan | X |  |  |  |  |  |  |  | X |  | X |
| MicrobeAnnotator | X | X |  |  |  |  |  |  |  |  | X |
| EggNOG-Mapper | X | X | X | X |  |  |  |  | X | X |  |

### Modes for running MetaCerberus

MetaCerberus has three basic modes: 1) quality control (QC) for raw reads, 2) formatting/gene prediction, and 3) annotation (**Fig 1**). MetaCerberus can use three different input files: 1) raw read data from any sequencing platform (Illumina, PacBio, or Oxford Nanopore), 2) assembled contigs, as MAGs, vMAGs, isolate genomes, or a collection of contigs, 3) amino acid fasta (.faa), previously called pORFs (**Fig 1**). We offer customization, including running databases all together, individually or specifying select databases. For example, if a user wants to run a prokaryotic or eukaryotic-specific KOfam, or an individual database alone such as dbCAN, both are easily customized within MetaCerberus. In future versions, we will provide viral and phage-specific KO modules to run individually. In QC mode, raw reads are quality controlled via fastqc (version v0.12.1) prior and post trim (**43**). Raw reads are then trimmed via data type; if the data is Illumina or PacBio, fastp (version 0.23.4) is called, otherwise it assumes the data is Oxford nanopore then Porechop (version v0.2.4) is utilized (**43-45, Fig 1**). Post quality-control trimmed reads are converted to fasta without quality (**Fig 1**). If Illumina reads are utilized, an optional bbmap (version 39.01) step to remove the phiX174 genome is available. Phage phiX174 is a common contaminant within the Illumina platform as it is their library spike-in control (**46-47**). We highly recommend this removal if viral analysis is conducted, as it would provide false positives to ssDNA microviruses within the sample.

**Figure 1:**
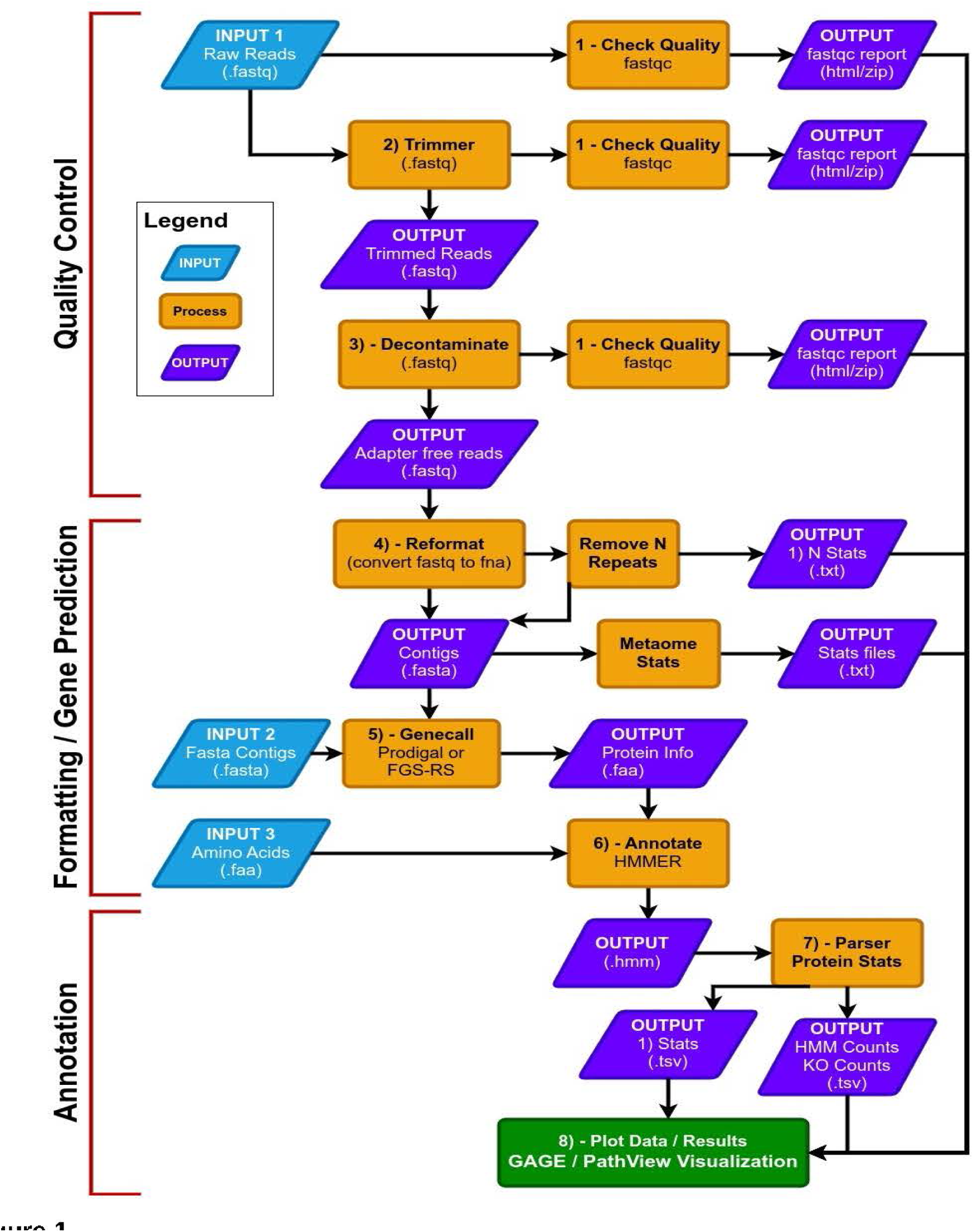
Flowgraph of the MetaCerberus pipeline. MetaCerberus has three input data formats which include raw reads, contigs, or previously called pORFs. Quality control step is mainly utilized for raw reads only. Contigs have a formatting mode were they are quality controlled for N presence, followed by N removal, and also provides basic contig statistics using Metaome Stats (e.g., N50, N90 etc). Gene calling currently offers prodigal or FragGeneScanRs for pORF calling. Gene prediction is completed with HMMER/HMM against KEGG and FOAM KOs (all by default). Users can select additional databases such as CAZy, COG, PHROG, VOG, and pVOGs for viruses, or selective KOfams for prokaryotes or eukaryotes. With running four or more samples it provides a PCA for KEGG and FOAM KOs, a basic run metric dashboard, as well as differential statistics using DESeq2/edgeR, and pathway enrichment using GAGE R followed by plotting in Pathview R.

In the formatting and gene prediction mode, contigs and genomes are checked for N repeats. These N repeats are removed by default. We impute contig/genome statistics (e.g., N50, N90, max contig) via our custom module Metaome Stats. Contigs are converted to pORFs via Prodigal or FragGeneScanRs as specified by user preference (**48, Fig 1**). Scaffold annotation is not recommended due to N’s providing ambiguous annotation. Both callers can be used via our ––super option, and we recommend using FragGeneScanRs for samples rich in eukaryotes as it performed better in our hands than Prodigal (unpublished data). HMMER searches against the above databases via user specified bitscore and e-values or our minimum defaults (i.e., bitscore = 25, e-value = 1 x 10^-9^).

There are six general rules followed by MetaCerberus for functional annotation. Rule 1 is the *score pre-filtering module* for pORFs thresholds: each pORF match to an HMM is recorded by default or user-selected e-value/bit scores per database independently, across all databases, or per user specification of the selected database. Rule 2 is imputed for *non-overlapping dual domain module* pORF threshold: if two HMM hits are non-overlapping from the same database, both are counted as long as they are the within the default or user selected e-value/bit scores. Rule 3 is computed as the *winner take all module* for overlapping pORFs: if two HMM hits are overlapping (>10 amino acids) from the same database the lowest resulting e-value and highest bit score wins. Rule 4 is *similar match independent accession module* for a single pORF: if both hits within the same database have equal values for both e-value and bit score but are different accessions from the same database (e.g., KO1 and KO3) then both are reported. Rule 5 is the *whole count incomplete exclusion module* filter only allows whole discrete integer counting. Rule 6 the *dual independent overlapping domain module* for convergent binary domain pORFs. If two domains within a pORF are overlapping <10 amino acids (e.g, COG1 and COG4) then both domains are counts and reported due to the dual domain issue within a single pORF. If a function hits multiple pathways within an accession, both are counted, in pathway roll-up, as many proteins function in multiple pathways.

### Statistics and visualization

MetaCerberus, as previously mentioned, provides genome and contig statistics via MetaOme stats; it also offers seamless integration into automatic differential statistics, visualizations, pathway enrichment, and pathway integration viewing. DESeq2 and edgeR negative binomial distribution differential statistic tools are available to users by selection (default is DESeq2) (**49-50**). The outputs from DESeq2, edgeR, or both are automatically enriched for pathway analysis in GAGE (Generally Applicable Gene-set Enrichment for Pathway Analysis) R (**51**). GAGE outputs are loaded into Pathview R to visualize differential pathways across user-specified experiments (**52**). These outputs include differential KEGG heatmaps from Pathview, volcano plots, and gene level heatmaps (**Fig S1**).

A sample dashboard visualization is provided for all data input types (e.g., reads, contigs and/or genomes) with a number of pORF called, MetaOme stats (i.e., genome statistics, N50, N90, etc., for genomes/contigs only), PCA with sample sets of >3, and the number of annotated hits for all databases or user select specifications (**Fig S1**).

### Across tool comparisons

Tools compared across MetaCerberus (version 1.1) include DRAM (version 1.4.6), InterProScan (version 5.60-92.0), EggNog-Mapper (version 2.1.8), MicrobeAnnotator (version 2.0.5), and PROKKA (version 1.1). All comparisons were completed on a Dual 8-Core Intel Xeon E5-2667 CPU @ 3.2GHz (16 cores) using 128GB RAM. MPP testing of MetaCerberus was completed on five nodes of a Dual 18-Core Intel Xeon Gold 6154 CPU @ 3.00GHz (36 cores/node). All genomes used in our study are available at https://osf.io/3uz2j/. For further testing of MetaCerberus, we used five distinct genospecies, *Rhizobium leguminosarum,* against five distinct *Exiguobacterium* spp. available at https://github.com/raw-lab/MetaCerberus/tree/main/data/rhizobium_test (**Table S1**). Viruses from permafrost that were used in the DRAM paper (https://www.ncbi.nlm.nih.gov/nuccore/QGNH00000000) were compared directly to MetaCerberus and DRAM (**19**).

### Data availability

Sequence files, genome files, and supplemental data are available at https://osf.io/3uz2j/. Databases are also freely available at https://osf.io/3uz2j/. All code is available at www.github.com/raw-lab/metacerberus.

### Contributing to MetaCerberus and Fungene

MetaCerberus is a community resource as is recently acquired FunGene (http://fungene.cme.msu.edu/). We welcome contributions of other experts expanding annotation of all domains of life (viruses, phages, bacteria, archaea, eukaryotes).

Please send us an issue on our MetaCerberus GitHub. (www.github.com/raw-lab/metacerberus/issue); we will fully annotate your genome, add suggested pathways/metabolisms of interest, make custom HMMs to be added to MetaCerberus and FunGene.

## Results

### Database size and download time

Formatting and downloading are required steps in functional annotation and both depend on database size. Substantial databases take up large amounts of costly disk space and require expensive computers with large amounts of costly RAM for analysis. MetaCerberus database size is 3.8 GB, with a download time of ∼4 mins, and database format time is zero because they are pre-formatted already for the user (**Table 2**). DRAM database download requires 710 GB of disc space, and requires ∼3 days to download completely (**Table 2**). According to the DRAM readme, KEGG Genes and UniRef90 need ∼500 GB of disc space and ∼512 GB of RAM to process the complete database (**19,** https://github.com/WrightonLabCSU/DRAM). This database size difference is due UniRef90 updates since their original release in 2020. DRAM can run with more processors within a single node but is not set up for multi-node like MetaCerberus. The InterProScan database is 14 GB, which took ∼2.45 h to install (**Table 2**). PROKKA had the smallest database at 636 MB and had the fastest install of ∼3 ½ minutes (**Table 2**). MicrobeAnnotator requires at least ∼237 GB for its full version and ∼0.65 GB for its light version (**Table 2**, **22**).

**Table 2:** Database size, download, and formatting time across tools.

| Tool | Time | Disk | Version |
| --- | --- | --- | --- |
| DRAM | ~ 3 days | ~710GB | v1.4.6 |
| InterProScan | ~ 2:45:59.23 | 14GB | v5.60-92.0 |
| Metacerberus | ~ 0:04:14.29 | 3.8GB | v1.1 |
| PROKKA | ~ 0:03:28.68 | 607M | v1.14.6 |
| EggNOG-Mapper | ~14:33:31.74 | 31GB | V2.1.8 |
| MicrobeAnnotator | >3 days | ~237GB | v2.0.5 |

### Computational resource comparison

We compared MetaCerberus to DRAM, InterProScan, and PROKKA for the time used per genome, RAM utilization, and disk space used across 100 randomly selected bacterial genomes within GTDB (**Table S1, Fig 2**). Generally, PROKKA had the highest processing speed per genome (∼48 sec median, **Fig 2**). InterProScan had the slowest at ∼21 min per genome median time (**Fig 2**). DRAM was ∼5 mins faster per genome than MetaCerberus (i.e., 10 mins vs 15 mins) (**Fig 2**). MetaCerberus and PROKKA had the lowest RAM, followed by InterProScan (**Fig 2**). DRAM using default parameters had the highest RAM observed (**Fig 2**). DRAM had the lowest disc space due to the deletion of files post-finalization, with PROKKA having the most disc space (**Fig 2**). EggNOG-mapper using HMMs was initially compared to MetaCerberus; however, further testing was not completed due to the high run time failure rate. On average, EggNOG-mapper failed to finish annotation 32% of the time using 148 randomly selected GTDB bacterial/archaeal genomes used by other tools (**Table S3**). Approximately 16% of the genomes failed after running for two days; another 16% could not annotate even when other tools, including MetaCerberus, had no issue (**Table S3**). The average annotation time with EggNOG-mapper v2 was ∼53 min, with a median of ∼30 min (**Table S3**). It was the slowest tool tested and thus removed from further comparisons. MicrobeAnnotator didn’t functionally install the database correctly. We could not successfully use the code; thus, it was removed from further comparisons..

**Figure 2:**
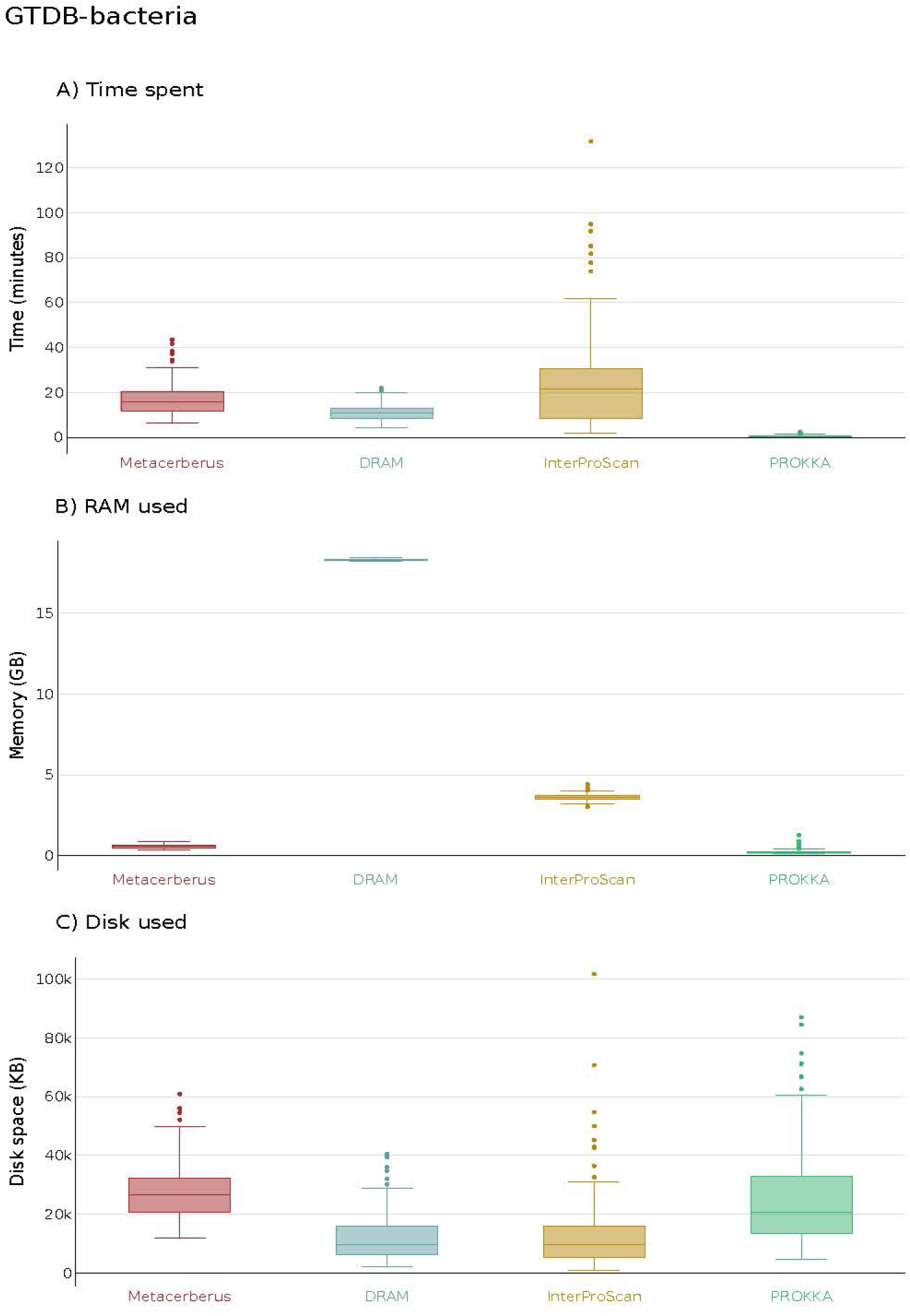
Computational resource comparison. DRAM, InterProScan, PROKKA, and MetaCerberus are compared computationally for time to complete each genome annotation, RAM required to complete annotation per genome, and disc space needed for inputs/outputs. The 100 randomly selected bacterial genomes were from GTDB (**Table S1**).

### Automatic statistical and pathway analysis

MetaCerberus provides automatic differential statistics, pathway gene enrichments, and KEGG map-based heatmaps in Pathview R for data exploration, data mining, and hypothesis generation. As a test, we compared five distinct genospecies, *Rhizobium leguminosarum,* against five distinct *Exiguobacterium* spp. using MetaCerberus using both DESeq2 and edgeR (**Table S3**). These genomes were selected as a comparison due to differential pathways within the comparison genomes. Rhizobium are symbiotic nitrogen fixers containing both nitrogenase for nitrogen fixation and nodulation genes for symbiotic nodule formation within legume roots (**53**). The *Exiguobacterium* spp. Have a bright orange colony morphology color from biosynthetically made carotenoids; it is hypothesized that the carotenoid diaponeurosporene-4-oic acid from the C_30_ carotenoid biosynthesis pathway is what provides the distinctive orange color (**54**). Chemical studies have suggested other carotenoids are present within *Exiguobacterium spp*. that contribute to the orange colony color (**55**). MetaCerberus found differential pathway assignments using DESeq2 and Pathview for carotenoid biosynthesis, ABC transporters, and phosphotransferase system (including nitrogen regulation) (**Fig S2-4**). edgeR found an additional pathway in benzoate degradation that wasn’t found in DESeq2 (**Fig S5**).

### Annotation comparisons

PROKKA, DRAM, and MetaCerberus all use Prodigal for pORF calling. MetaCerberus also provides an extra pORF caller FragGeneScanRs. InterProScan uses the EMBOSS getorf pORF caller, which in all cases had lower pORFs than Prodigal regardless of the genome kingdom type (e.g., bacteria, archaea, CPR, phage, archaeal virus or eukaryotic virus) (**Fig S6**). Generally, PROKKA, DRAM, and MetaCerberus had similar pORF calling numbers; however, DRAM did call more pORF from eukaryotic viruses (**Fig S6**).

Furthermore, we compared MetaCerberus to DRAM, InterProScan, and PROKKA for whether a pORF was annotated, listed as hypothetical, or unknown (no annotation). We randomly selected 100 unique bacterial genomes from GTDB, 100 unique archaea genomes from GTDB, 100 unique phage genomes from INPHARED, 100 unique eukaryotic viral genomes from RefSeq, 78 CPR genomes, and 82 archaeal viral genomes for these annotation tests (**Table S1**). MetaCerberus, DRAM, and InterProScan protein modes had similar annotation results of ∼78-83% for bacteria, with InterProScan being the highest at 83% (**Fig 3**). InterProScan using nucleotide mode had the lowest annotation amount across all kingdoms (**Fig 3-4**). PROKKA had ∼50% of the pORFs as annotated and hypothetical for bacteria and ∼60% hypothetical for archaea (**Fig 3**). CPR annotation InterProScan had the highest at 70%, followed by DRAM at 66%, then MetaCerberus at 61% (**Fig 3**). MetaCerberus and PROKKA had fewer unknowns than DRAM for bacteria, archaea, and CPR genomes (**Fig 3**). PROKKA annotated very few CPR pORFs, with the majority >60% being hypothetical proteins (**Fig 3**). DRAM generally doesn’t find many hypothetical proteins or lists them as unknown across domains of the tree of life.

**Figure 3:**
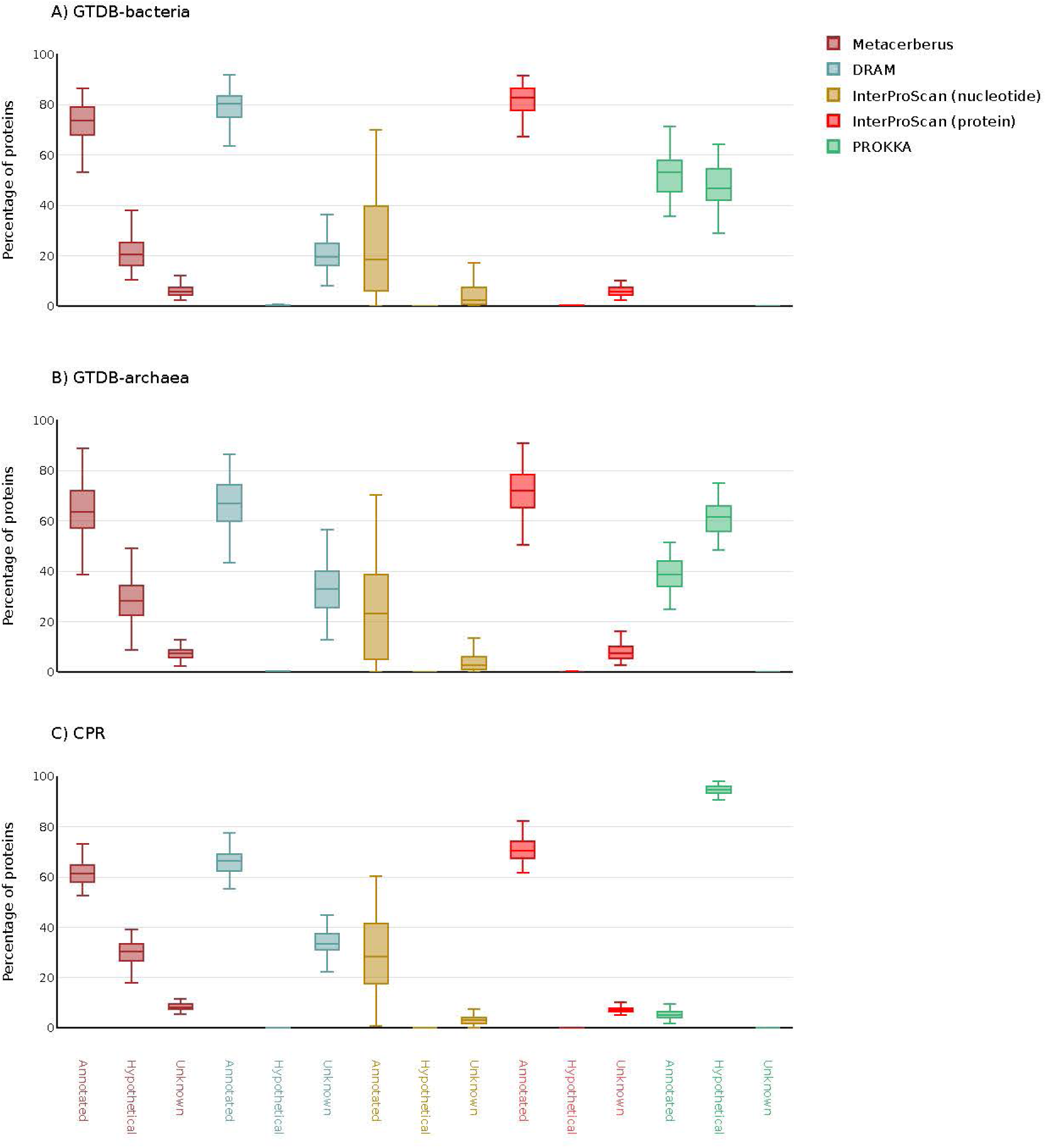
Annotation comparison across cellular domains of life (bacteria, archaea, CPR). MetaCerberus was compared to DRAM, InterProScan, and PROKKA for annotation across various genomes. Supplemental materials include the genomes for bacteria, archaea, and CPR used in this comparison (**Table S1**).

**Figure 4:**
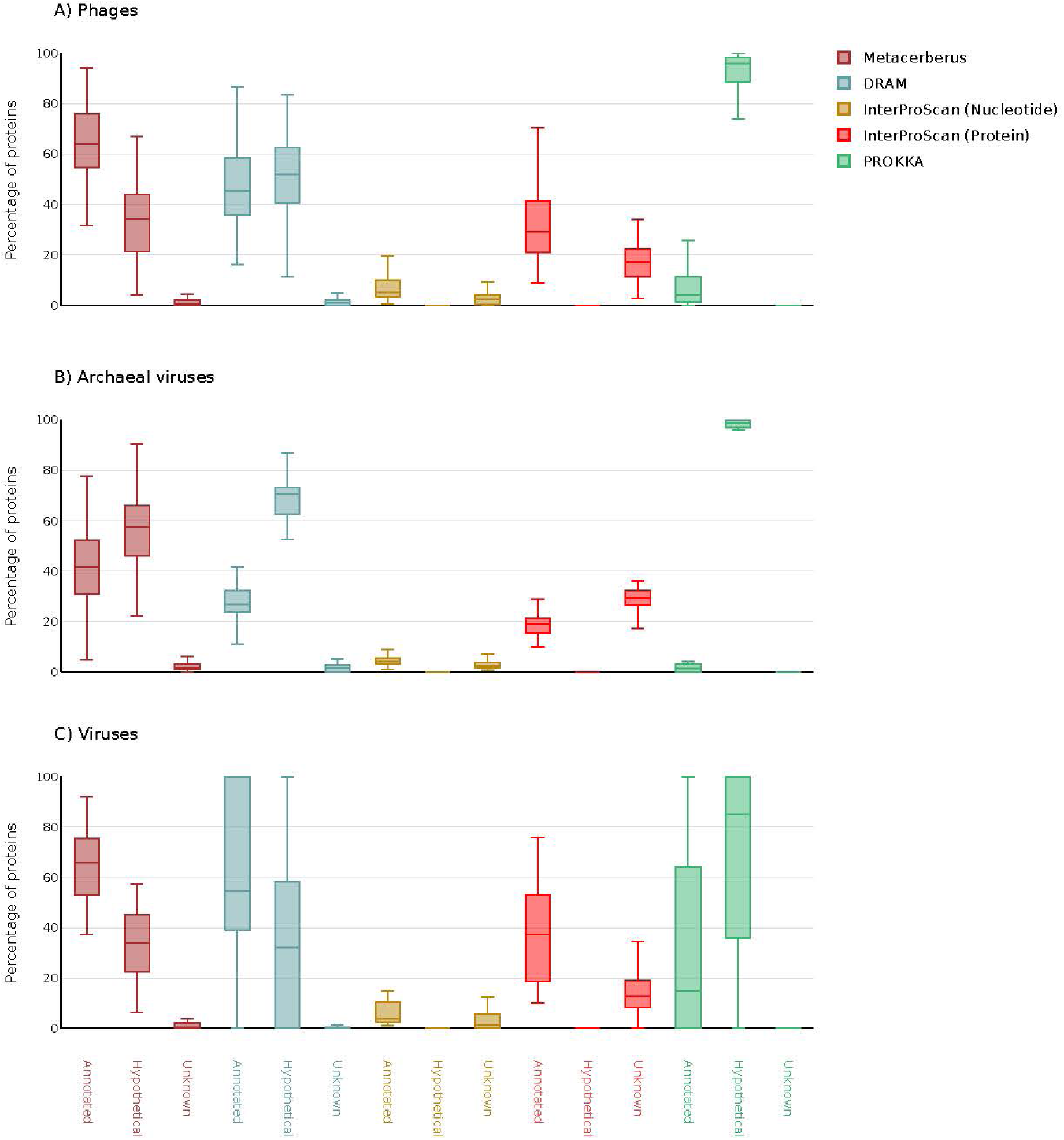
Annotation comparison across viruses infecting differential cellular domains (phage, archaeal viruses, eukaryotic viruses). MetaCerberus was compared to DRAM, InterProScan, and PROKKA for annotation across various genomes. The genomes are listed for phage, archaeal viruses, and eukaryotic viruses in supplemental materials (**Table S1**).

MetaCerberus performs better for viruses, phages, and archaeal viruses (**Fig 4**). MetaCerberus annotates more per genome >63 % phages, >65 % viruses, and >41% archaeal viruses based on median values (**Fig 4)**. MetaCerberus outperforms across all viruses (e.g., phages, eukaryotic viruses, and archael viruses) providing more annotations, fewer hypotheticals, and fewer unknowns compared to DRAM, InterProScan, and PROKKA (**Fig 4)**. MetaCerberus annotates more KOs from KOfams than DRAM across all domains (**Fig 5**). PROKKA and InterProScan don’t provide KOs; therefore, we couldn’t compare KOs found across domains to MetaCerberus.

**Figure 5.**
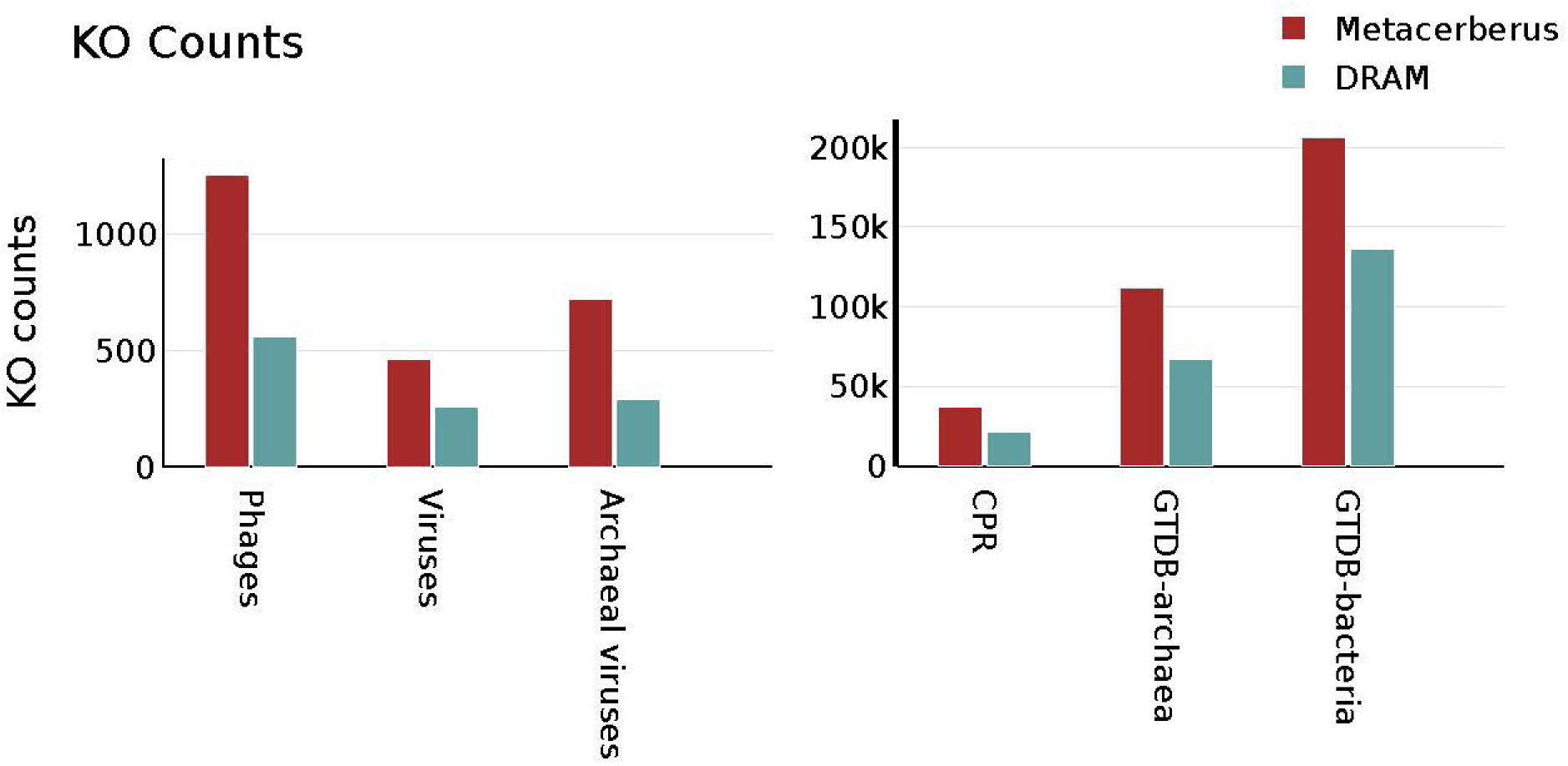
DRAM vs. MetaCerberus KO annotation comparison across the domains of life. DRAM and MetaCerberus utilize KOfams for KEGG KO assignment if the user doesn’t provide a KEGG KO database separately. The genomes for the comparison are listed in supplemental materials (**Table S1**). The e-values and bitscore can vary between DRAM and MetaCerberus. In this comparison, we choose the default dbCAN e-value option of <1e^-15^ and the default bitscore of 60 for DRAM.

To better compare to DRAM-v, the only other tool exclusively for viruses and phages, we analyzed a virome containing 1,907 viral populations (VPs) obtained from Swedish permafrost. Based on time, MetaCerberus took 99 mins to complete the annotation compared to 141.75 mins for DRAM (**Fig 6**). When MetaCerberus is utilized at it’s full potential with RAY it only takes 12.5 mins for the same dataset (**Fig 6**). RAM was significant less with MetaCerberus vs. DRAM-v, both in MPP and non-MPP mode, with <500 Mb of RAM compared to 18.7 Gb with DRAM-v (**Fig 6**). MetaCerberus had more annotations than DRAM-v for the Swedish permafrost virome across shared databases (i.e., KO, CAZy, and VOG) (**Fig S7**).

**Figure 6.**
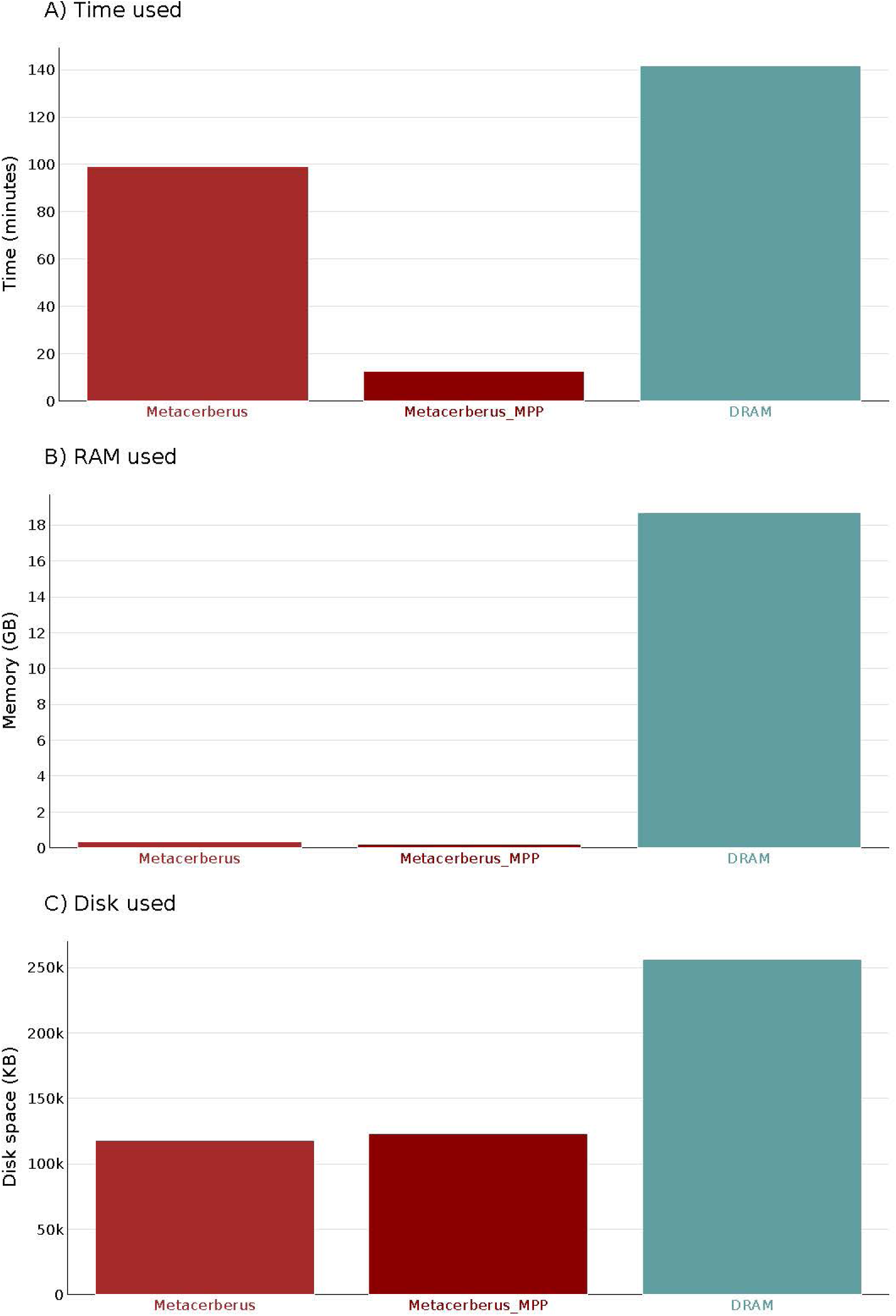
DRAM vs. MetaCerberus computational resource comparison. A virome from Swedish permafrost containing 1,907 VPs were compared computationally for time to complete annotation, RAM required to complete annotation, and disc space needed for inputs/outputs. MPP testing for MetaCerberus utilized five nodes for comparisons.

## Discussion

MetaCerberus provides a low memory, robust, scalable, and rapid annotation across the tree of life, exclusively using HMMs/HMMER. HMMER is a powerful tool to find pORFs that may be missed by standard homology-based tools due to its protein domain centric and supervised machine-learning nature. It’s rarely used alone due to the speed and time required to finish annotation. MetaCerberus has provided a solution to this scaling issue using RAY and algorithms needed from MerCat2. EggNOG-Mapper v2 is the only other tool that exclusively provides HMMs/HMMER-based annotation alone. MetaCerberus runs twice as fast on a single node than EggNOG-Mapper v2 without RAY. MetaCerberus with RAY is three times as fast as EggNOG-Mapper v2 (data not shown) on a database that is 1/3 smaller.

Generally, MetaCerberus performs better for viral and phage annotation when directly compared to DRAM-v. DRAM finds more pORFs than MetaCerberus (**Fig S6**) due to it using the –meta option in Prodigal for viruses; whereas, MetaCerberus uses a standard for complete genomes in this case but still can annotate viral genomes better than DRAM on a much smaller database. Viruses, archaeal viruses, and phages are a grand challenge to unlock the ‘unknown’ and ‘hypothetical’ functions within their genomes.

As data scales, computational time, memory, and waiting for results will take longer. Scalable tools like MetaCerberus are needed as we approach Petabyte levels of sequencing. MetaCerberus provides a further community resource to annotate the unknowns of our biosphere. Lastly, MetaCerberus provides a robust tool kit to annotate the entire tree of life at scale.

## Acknowledgments

We also acknowledge the University Research Computing and the College of Computing and Informatics for computational and logistical support. We must further acknowledge J Peter W Young at the University of York for our discussions on the nature of *Rhizobium* genomes, especially genospecies. For his insightful grammar edits and proofreading of multiple drafts, we acknowledge Bryan W. Fulghum.

## Supplemental Materials

**Table S1**: List of genomes used for computational comparisons. This includes randomly selected GTDB genomes for archaea and bacteria domains, phage genomes, archaeal viral genomes, CPR genomes, and RefSeq viral genomes.

**Figure S1**: Standard output dashboard for MetaCerberus. Outputs include a complete html drawn in plotly. Also, for comparisons of >3 genomes, FOAM and KEGG-based KO PCA are included.

**Figure S2**: Comparisons *Rhizobium* vs. *Exiguobacterium* genomes using MetaCerberus for carotenoid pathways in Pathview. KO counts from KEGG were normalized with DESeq2, enriched with GAGE, then plotted with Pathview R. Genomes are listed in supplemental materials (**Table S1**).

**Figure S3**: Comparisons *Rhizobium* vs. *Exiguobacterium* genomes using MetaCerberus for ABC transporters in Pathview. KO counts from KEGG were normalized with DESeq2, enriched with GAGE, then plotted with Pathview R. Genomes are listed in supplemental materials (**Table S1**).

**Figure S4**: Comparisons *Rhizobium* vs. *Exiguobacterium* genomes using MetaCerberus for phosphotransferase system in Pathview. KO counts from KEGG were normalized with DESeq2, enriched with GAGE, then plotted with Pathview R. Genomes are listed in supplemental materials (**Table S1**).

**Figure S5**: Comparisons *Rhizobium* vs. *Exiguobacterium* genomes using MetaCerberus for Benzoate degradation in Pathview. KO counts from KEGG was normalized with edgeR, enriched with GAGE, then plotted with Pathview R. Genomes are listed in supplemental materials (**Table S1**).

**Figure S6**: Comparisons of protein-coding open reading frame (pORF) calling across computational tools. All tools but InterProScan use Prodigal for pORFs. InterProScan uses the EMBOSS getorf tool.

**Figure S7**: Comparing DRAM-v vs. MetaCerberus for annotation across shared databases (i.e., KO, VOG, CAZy). The Swedish virome was utilized for comparisons.

## References

1. Illumina throughput specs (date accessed July 17^th^, 2023). https://www.illumina.com/systems/sequencing-platforms/novaseq-x-plus.html

2. Oxford throughput specs (date accessed July 17^th^, 2023). https://nanoporetech.com/about-us/news/highest-throughput-yet-promethion-breaks-7-terabase-mark

3. Genome Taxonomy Database (GTDB) release statistics (date accessed July 17^th^, 2023). https://gtdb.ecogenomic.org/

4. Short Read Archive Biosample Metagenomes (date accessed July 17^th^, 2023). https://www.ncbi.nlm.nih.gov/sra/?term=metagenomes

5. Bowers RM, Kyrpides NC, Stepanauskas R, Harmon-Smith M, Doud D, Reddy TBK, Schulz F, Jarett J, Rivers AR, Eloe-Fadrosh EA, Tringe SG, Ivanova NN, Copeland A, Clum A, Becraft ED, Malmstrom RR, Birren B, Podar M, Bork P, Weinstock GM, Garrity GM, Dodsworth JA, Yooseph S, Sutton G, Glockner FO, Gilbert JA, Nelson WC, Hallam SJ, Jungbluth SP, Ettema TJG, Tighe S, Konstantinidis KT, Liu WT, Baker BJ, Rattei T, Eisen JA, Hedlund B, McMahon KD, Fierer N, Knight R, Finn R, Cochrane G, Karsch-Mizrachi I, Tyson GW, Rinke C, Genome Standards C, Lapidus A, Meyer F, Yilmaz P, Parks DH, Eren AM, Schriml L, Banfield JF, Hugenholtz P, Woyke T. Minimum information about a single amplified genome (MISAG) and a metagenome-assembled genome (MIMAG) of bacteria and archaea. Nat Biotechnol. 2017;35(8):725–731.

6. Roux S, Adriaenssens EM, Dutilh BE, Koonin EV, Kropinski AM, Krupovic M, Kuhn JH, Lavigne R, Brister JR, Varsani A, Amid C, Aziz RK, Bordenstein SR, Bork P, Breitbart M, Cochrane GR, Daly RA, Desnues C, Duhaime MB, Emerson JB, Enault F, Fuhrman JA, Hingamp P, Hugenholtz P, Hurwitz BL, Ivanova NN, Labonté JM, Lee KB, Malmstrom RR, Martinez-Garcia M, Mizrachi IK, Ogata H, Páez-Espino D, Petit MA, Putonti C, Rattei T, Reyes A, Rodriguez-Valera F, Rosario K, Schriml L, Schulz F, Steward GF, Sullivan MB, Sunagawa S, Suttle CA, Temperton B, Tringe SG, Thurber RV, Webster NS, Whiteson KL, Wilhelm SW, Wommack KE, Woyke T, Wrighton KC, Yilmaz P, Yoshida T, Young MJ, Yutin N, Allen LZ, Kyrpides NC, Eloe-Fadrosh EA. Minimum Information about an Uncultivated Virus Genome (MIUViG). Nat Biotechnol. 2019 37(1):29–37.

7. Kieft K, Adams A, Salamzade R, Kalan L, Anantharaman K. vRhyme enables binning of viral genomes from metagenomes. Nucleic Acids Res. 2022;50(14):e83.

8. Hyatt D, Chen G-L, LoCascio PF, Land ML, Larimer FW, Hauser LJ. Prodigal: prokaryotic gene recognition and translation initiation site identification. BMC Bioinformatics. 2010;11:119.

9. Van der Jeugt F, Dawyndt P, Mesuere B. FragGeneScanRs: faster gene prediction for short reads. BMC Bioinformatics. 2022 3(1):198.

10. Emboss getorfs (date accessed July 17^th^, 2023). https://emboss.sourceforge.net/apps/cvs/emboss/apps/getorf.html

11. Besemer J, Borodovsky M. GeneMark: web software for gene finding in prokaryotes, eukaryotes and viruses. Nucleic Acids Res. 2005 33:W451–4.

12. Camacho C, Coulouris G, Avagyan V, Ma N, Papadopoulos J, Bealer K, Madden TL. BLAST+: architecture and applications. BMC Bioinform. 2009;10:421.

13. Steinegger M, Söding J. MMseqs2 enables sensitive protein sequence searching for the analysis of massive data sets. Nat Biotechnol. 2017 35(11):1026–1028.

14. Buchfink B, Xie C, Huson DH. Fast and sensitive protein alignment using DIAMOND. Nat Methods. 2015, 12(1):59–60.

15. O’Leary NA, Wright MW, Brister JR, Ciufo S, Haddad D, McVeigh R, Rajput B, Robbertse B, Smith-White B, Ako-Adjei D, Astashyn A, Badretdin A, Bao Y, Blinkova O, Brover V, Chetvernin V, Choi J, Cox E, Ermolaeva O, Farrell CM, Goldfarb T, Gupta T, Haft D, Hatcher E, Hlavina W, Joardar VS, Kodali VK, Li W, Maglott D, Masterson P, McGarvey KM, Murphy MR, O’Neill K, Pujar S, Rangwala SH, Rausch D, Riddick LD, Schoch C, Shkeda A, Storz SS, Sun H, Thibaud-Nissen F, Tolstoy I, Tully RE, Vatsan AR, Wallin C, Webb D, Wu W, Landrum MJ, Kimchi A, Tatusova T, DiCuccio M, Kitts P, Murphy TD, Pruitt KD. Reference sequence (RefSeq) database at NCBI: current status, taxonomic expansion, and functional annotation. Nucleic Acids Res. 2016 44(D1):D733–45.

16. UniProt Consortium. UniProt: the Universal Protein Knowledgebase in 2023. Nucleic Acids Res. 2023 51(D1):D523–D531.

17. Kanehisa M, Furumichi M, Tanabe M, Sato Y, Morishima K. KEGG: new perspectives on genomes, pathways, diseases and drugs. Nucleic Acids Res. 2017;45:D353–61.

18. Seemann T. Prokka: rapid prokaryotic genome annotation. Bioinformatics. 2014. 30(14):2068–9.

19. Shaffer M, Borton MA, McGivern BB, Zayed AA, La Rosa SL, Solden LM, Liu P, Narrowe AB, Rodríguez-Ramos J, Bolduc B, Gazitúa MC, Daly RA, Smith GJ, Vik DR, Pope PB, Sullivan MB, Roux S, Wrighton KC. DRAM for distilling microbial metabolism to automate the curation of microbiome function. Nucleic Acids Res. 2020 48(16):8883–8900.

20. Jones P, Binns D, Chang HY, Fraser M, Li W, McAnulla C, McWilliam H, Maslen J, Mitchell A, Nuka G, Pesseat S, Quinn AF, Sangrador-Vegas A, Scheremetjew M, Yong SY, Lopez R, Hunter S. InterProScan 5: genome-scale protein function classification. Bioinformatics. 2014 30(9):1236–40.

21. Cantalapiedra CP, Hernández-Plaza A, Letunic I, Bork P, Huerta-Cepas J. eggNOG-mapper v2: Functional Annotation, Orthology Assignments, and Domain Prediction at the Metagenomic Scale. Mol Biol Evol. 2021 38(12):5825–5829.

22. Ruiz-Perez CA, Conrad RE, Konstantinidis KT. MicrobeAnnotator: a user-friendly, comprehensive functional annotation pipeline for microbial genomes. BMC Bioinformatics. 2021 22(1):11.

23. Eddy SR. Accelerated Profile HMM Searches. PLoS Comput Biol. 2011 7(10):e1002195.

24. Hernández-Plaza A, Szklarczyk D, Botas J, Cantalapiedra CP, Giner-Lamia J, Mende DR, Kirsch R, Rattei T, Letunic I, Jensen LJ, Bork P, von Mering C, Huerta-Cepas J. eggNOG 6.0: enabling comparative genomics across 12 535 organisms. Nucleic Acids Res. 2023 51(D1):D389–D394.

25. Paysan-Lafosse T, Blum M, Chuguransky S, Grego T, Pinto BL, Salazar GA, Bileschi ML, Bork P, Bridge A, Colwell L, Gough J, Haft DH, Letunić I, Marchler-Bauer A, Mi H, Natale DA, Orengo CA, Pandurangan AP, Rivoire C, Sigrist CJA, Sillitoe I, Thanki N, Thomas PD, Tosatto SCE, Wu CH, Bateman A. InterPro in 2022. Nucleic Acids Res. 2023 January 6th;51(D1):D418–D427.

26. Mistry J, Chuguransky S, Williams L, Qureshi M, Salazar GA, Sonnhammer ELL, Tosatto SCE, Paladin L, Raj S, Richardson LJ, Finn RD, Bateman A. Pfam: The protein families database in 2021. Nucleic Acids Res. 2021 49(D1):D412–D419

27. Steinegger M, Meier M, Mirdita M, Vöhringer H, Haunsberger SJ, Söding J. HH-suite3 for fast remote homology detection and deep protein annotation. BMC Bioinformatics. 2019 20(1):473.

28. Fremin BJ, Bhatt AS, Kyrpides NC; Global Phage Small Open Reading Frame (GP-SmORF) Consortium. Thousands of small, novel genes are predicted in global phage genomes. Cell Rep. 2022 39(12):110984.

29. Jaffe AL, Castelle CJ, Matheus Carnevali PB, Gribaldo S, Banfield JF. The rise of diversity in metabolic platforms across the Candidate Phyla Radiation. BMC Biol. 2020 18(1):69.

30. Parks DH, Chuvochina M, Rinke C, Mussig AJ, Chaumeil PA, Hugenholtz P. GTDB: an ongoing census of bacterial and archaeal diversity through a phylogenetically consistent, rank normalized and complete genome-based taxonomy. Nucleic Acids Res. 2022 50(D1):D785–D794.

31. Virus Orthologous Groups (VOG) database https://vogdb.org/

32. Grazziotin AL, Koonin EV, Kristensen DM. Prokaryotic Virus Orthologous Groups (pVOGs): a resource for comparative genomics and protein family annotation. Nucleic Acids Res. 2017 January 4th;45(D1):D491–D498. doi: 10.1093/nar/gkw975. Epub 2016 October 26th. PMID: 27789703; PMCID: PMC5210652.

33. Camargo AP, Nayfach S, Chen IA, Palaniappan K, Ratner A, Chu K, Ritter SJ, Reddy TBK, Mukherjee S, Schulz F, Call L, Neches RY, Woyke T, Ivanova NN, Eloe-Fadrosh EA, Kyrpides NC, Roux S. IMG/VR v4: an expanded database of uncultivated virus genomes within a framework of extensive functional, taxonomic, and ecological metadata. Nucleic Acids Res. 2023 51(D1):D733–D743.

34. Cook R, Brown N, Redgwell T, Rihtman B, Barnes M, Clokie M, Stekel DJ, Hobman J, Jones MA, Millard A. INfrastructure for a PHAge REference Database: identification of large-scale biases in the current collection of cultured phage genomes. Phage (New Rochelle). 2021 2(4):214–223.

35. Terzian P, Olo Ndela E, Galiez C, Lossouarn J, Pérez Bucio RE, Mom R, Toussaint A, Petit MA, Enault F. PHROG: families of prokaryotic virus proteins clustered using remote homology. NAR Genom Bioinform. 2021 3(3):lqab067.

36. Figueroa III JL, Panyala A, Colby S, Friesen ML, Tiemann L, White III RA. MerCat2: a versatile k-mer counter and diversity estimator for database-independent property analysis obtained from omics data. bioRxiv, 2022.

37. Aramaki T, Blanc-Mathieu R, Endo H, Ohkubo K, Kanehisa M, Goto S, Ogata H. KofamKOALA: KEGG Ortholog assignment based on profile HMM and adaptive score threshold. Bioinformatics. 2020 36(7):2251–2252.

38. Prestat E, David MM, Hultman J, Taş N, Lamendella R, Dvornik J, Mackelprang R, Myrold DD, Jumpponen A, Tringe SG, Holman E, Mavromatis K, Jansson JK. FOAM (Functional Ontology Assignments for Metagenomes): a Hidden Markov Model (HMM) database with environmental focus. Nucleic Acids Res. 2014 42(19):e145.

39. Galperin MY, Wolf YI, Makarova KS, Vera Alvarez R, Landsman D, Koonin EV. COG database update: focus on microbial diversity, model organisms, and widespread pathogens. Nucleic Acids Res. 2021 49(D1):D274–D281.

40. Yin Y, Mao X, Yang J, Chen X, Mao F, Xu Y. dbCAN: a web resource for automated carbohydrate-active enzyme annotation. Nucleic Acids Res. 2012;40:W445–51.

41. Lombard V, Golaconda Ramulu H, Drula E, Coutinho PM, Henrissat B. The carbohydrate-active enzymes database (CAZy) in 2013. Nucleic Acids Res. 2014;42:D490–5.

42. Katoh K, Standley DM. MAFFT multiple sequence alignment software version 7: improvements in performance and usability. Mol Biol Evol. 2013 30(4):772–80.

43. FASTQC https://github.com/s-andrews/FastQC

44. Chen S, Zhou Y, Chen Y, Gu J. fastp: an ultra-fast all-in-one FASTQ preprocessor. Bioinformatics. 2018 34(17):i884–i890.

45. Porechop https://github.com/rrwick/Porechop

46. Moustafa A, Xie C, Kirkness E, Biggs W, Wong E, Turpaz Y, Bloom K, Delwart E, Nelson KE, Venter JC, Telenti A. The blood DNA virome in 8,000 humans. PLoS Pathog. 2017 13(3):e1006292.

47. Bushnell, Brian. BBMap: A Fast, Accurate, Splice-Aware Aligner. United States.

48. MetaOme Stats https://github.com/raw-lab/metaome_stats

49. Love MI, Huber W, Anders S. Moderated estimation of fold change and dispersion for RNA-seq data with DESeq2. Genome Biol. 2014 15(12):550.

50. Robinson MD, McCarthy DJ, Smyth GK. edgeR: a Bioconductor package for differential expression analysis of digital gene expression data. Bioinformatics. 2010 26(1):139–40.

51. Luo W, Friedman MS, Shedden K, Hankenson KD, Woolf PJ. GAGE: generally applicable gene set enrichment for pathway analysis. BMC Bioinformatics. 2009 10:161.

52. Luo W, Brouwer C. Pathview: an R/Bioconductor package for pathway-based data integration and visualization. Bioinformatics. 2013 29(14):1830–1.

53. Young JPW, Moeskjær S, Afonin A, Rahi P, Maluk M, James EK, Cavassim MIA, Rashid MH, Aserse AA, Perry BJ, Wang ET, Velázquez E, Andronov EE, Tampakaki A, Flores Félix JD, Rivas González R, Youseif SH, Lepetit M, Boivin S, Jorrin B, Kenicer GJ, Peix Á, Hynes MF, Ramírez-Bahena MH, Gulati A, Tian CF. Defining the *Rhizobium leguminosarum* Species Complex. Genes (Basel). 2021 12(1):111.

54. White RA 3rd, Soles SA, Gavelis G, Gosselin E, Slater GF, Lim DSS, Leander B, Suttle CA. The Complete Genome and Physiological Analysis of the Eurythermal Firmicute Exiguobacterium chiriqhucha Strain RW2 Isolated From a Freshwater Microbialite, Widely Adaptable to Broad Thermal, pH, and Salinity Ranges. Front Microbiol. 2019 9:3189.

55. Jinendiran S, Dahms HU, Dileep Kumar BS, Kumar Ponnusamy V, Sivakumar N. Diapolycopenedioic-acid-glucosyl ester and keto-myxocoxanthin glucoside ester: Novel carotenoids derived from *Exiguobacterium acetylicum* S01 and evaluation of their anticancer and anti-inflammatory activities. Bioorg Chem. 2020 103:104149.

