## Supplementary figures and images for "MetaCerberus: distributed highly parallelized scalable HMM-based implementation for robust functional annotation across the tree of life"

### Fig S1

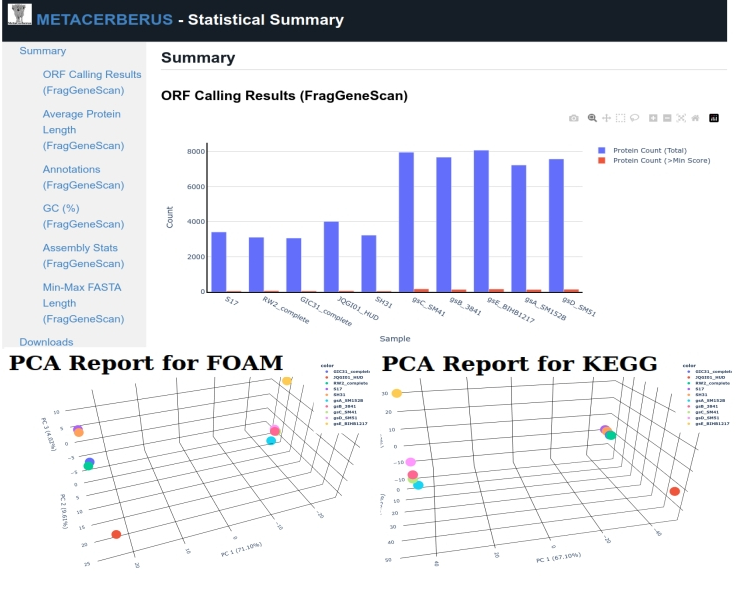

### Supplemental Fig S2-S7

#### Fig S2

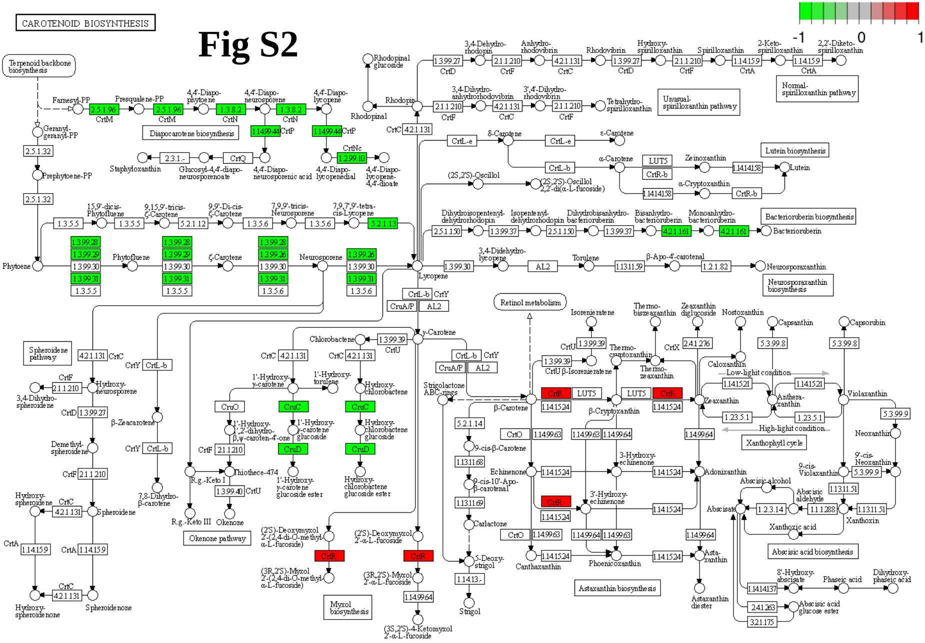


Data on KEGG graph
Rendered by Pathview

#### Fig S3

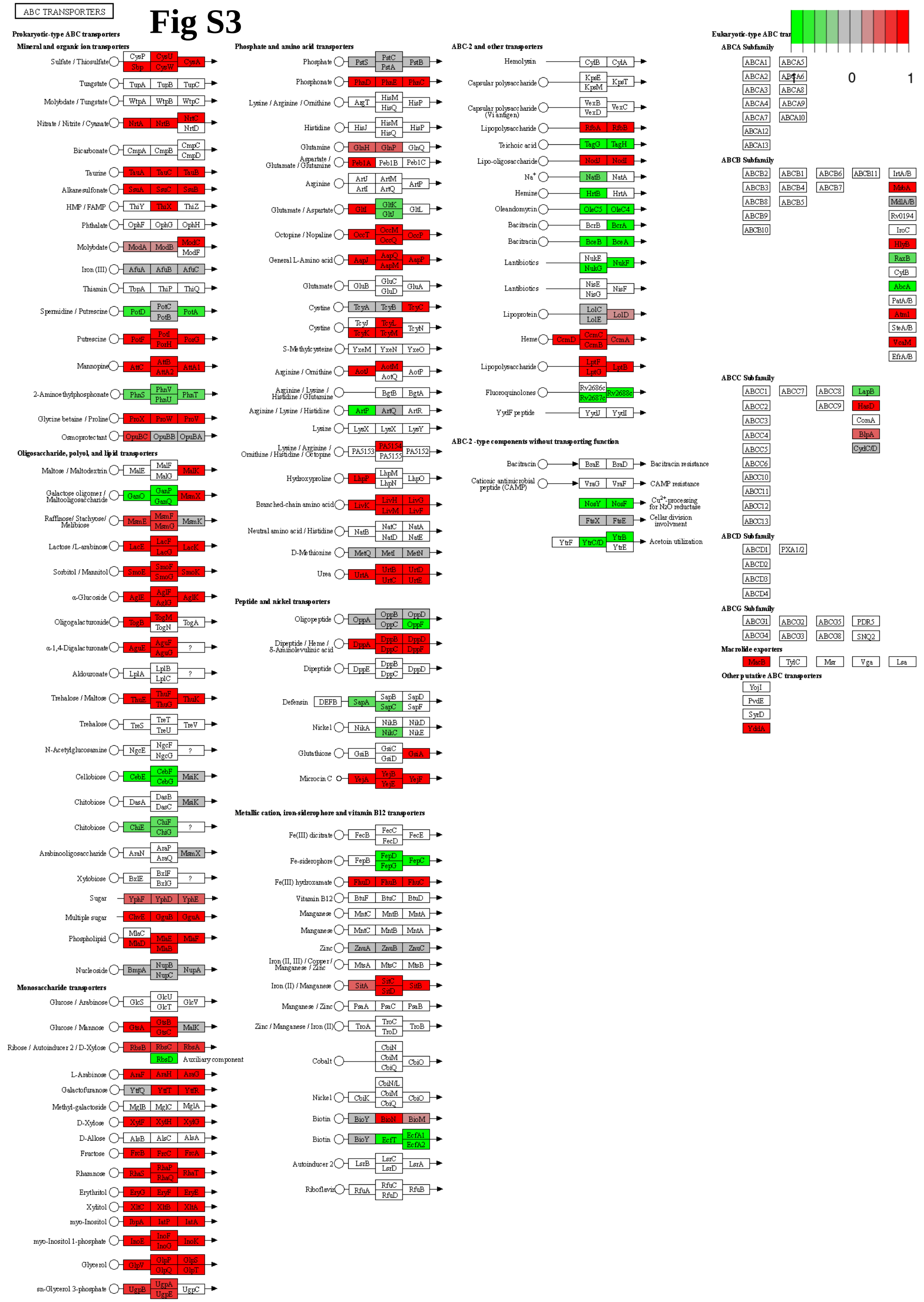


Data on KEGG graph
Rendered by Pathview

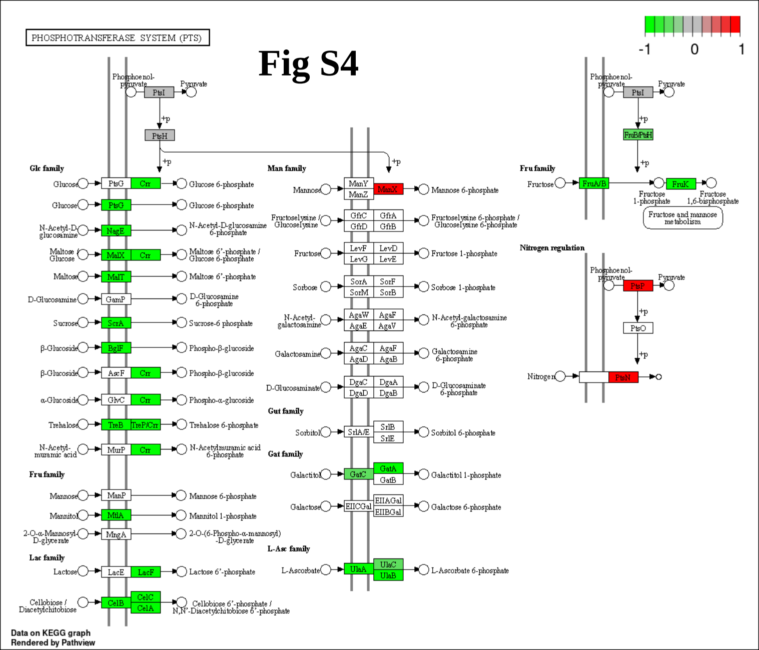

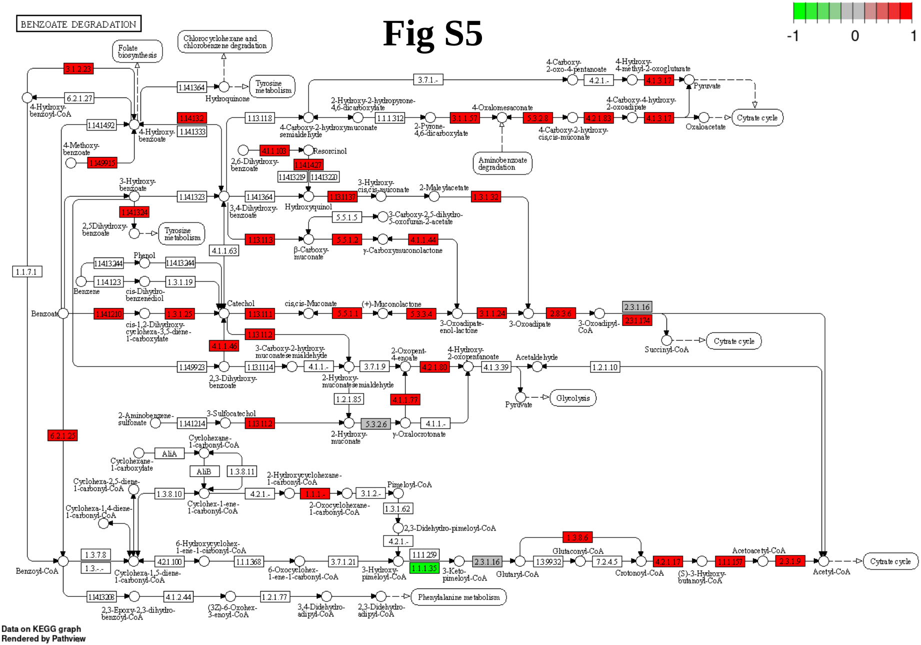

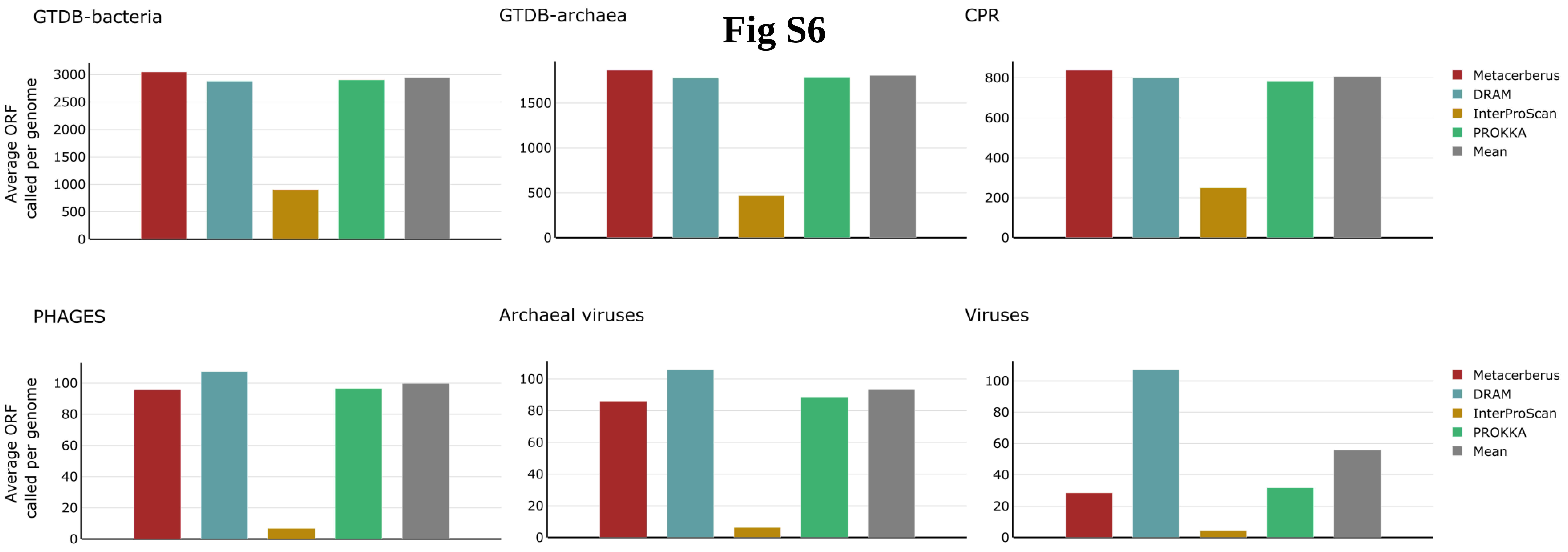

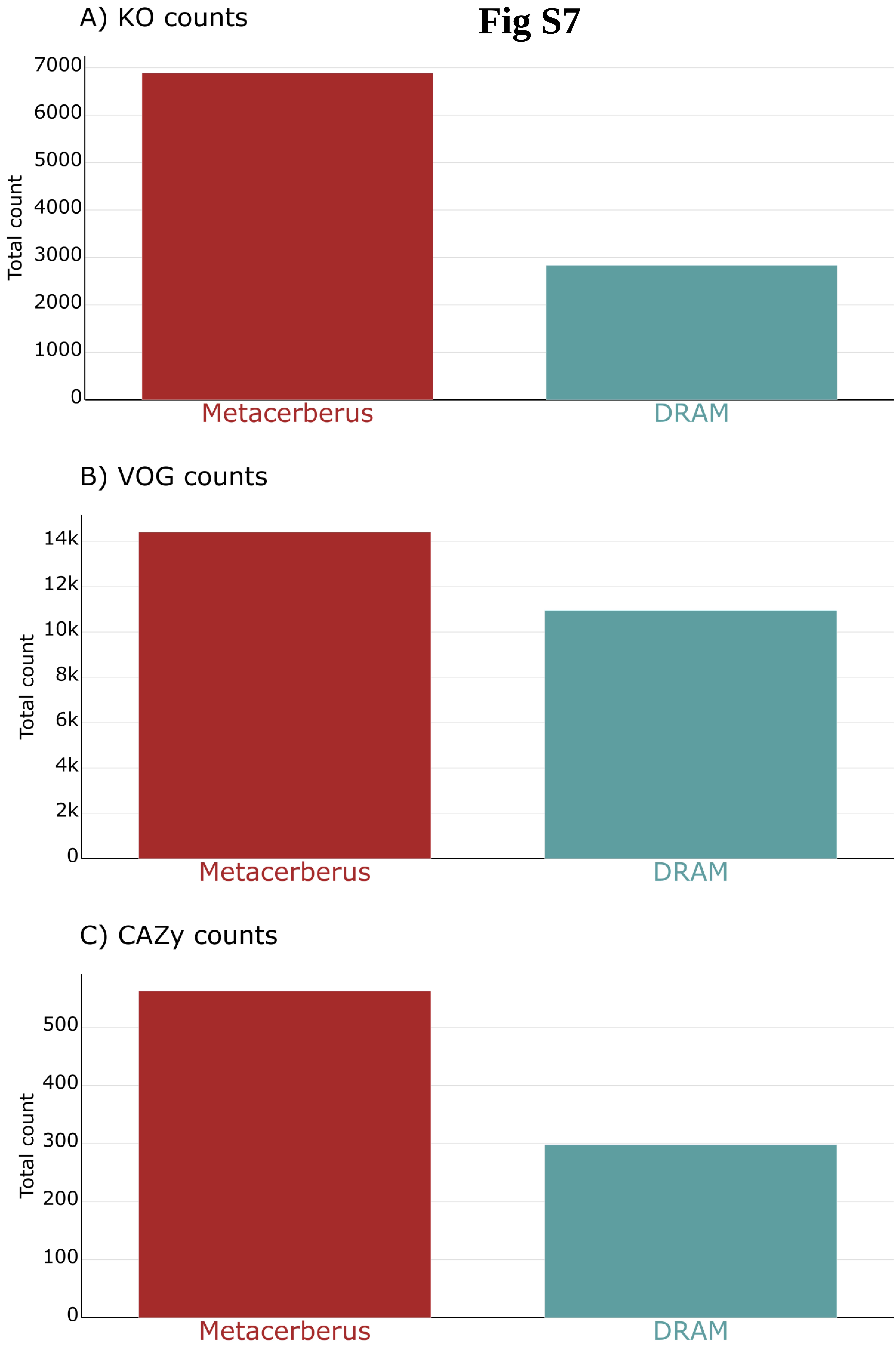
